# Leaf and flower consumption modulate the drinking behavior in a folivorous-frugivorous arboreal mammal

**DOI:** 10.1101/2020.07.20.211706

**Authors:** Óscar M. Chaves, Vanessa B. Fortes, Gabriela P. Hass, Renata B. Azevedo, Kathryn E. Stoner, Júlio César Bicca-Marques

## Abstract

Water is vital for the survival of any species because of its key role in most physiological processes. However, little is known about the non-food-related water sources exploited by arboreal mammals, the seasonality of their drinking behavior and its potential drivers (including diet composition, temperature, and rainfall). We investigated this subject in 14 wild groups of brown howler monkeys (*Alouatta guariba clamitans*) inhabiting small, medium, and large Atlantic Forest fragments in southern Brazil. We found a wide variation in the mean rate of drinking among groups (range=0-16 records/day). Streams (44% of 1,258 records) and treeholes (26%) were the major types of water sources, followed by bromeliads in the canopy (16%), pools (11%), and rivers (3%). The type of source influenced whether howlers used a hand to access the water or not. Drinking tended to be evenly distributed throughout the year, except for a slightly lower number of records in the spring than in the other seasons, but it was unevenly distributed during the day. It increased in the afternoon in all groups, particularly during temperature peaks around 15:00 and 17:00. We found via generalized linear mixed modelling that the daily frequency of drinking was mainly influenced by flower (negatively) and leaf (positively) consumption, whereas fruit consumption, fragment size, rainfall, and mean ambient temperature played negligible roles. The influence of leaf consumption is compatible with the ‘metabolite detoxification hypothesis,’ which states that the processing of this fibrous food requires the ingestion of larger volumes of water to help in the detoxification/excretion of its metabolites. In sum, we found that irrespective of habitat size and climatic conditions, brown howlers seem to seek a positive water balance by complementing preformed and metabolic water with drinking water, even when it is associated with a high predation risk in terrestrial sources.

## Introduction

Water is an essential chemical substance for all animals, not only because it represents a large percentage of whole-body mass, but because it is the medium within which the chemical reactions and physiological processes of the body take place [1–3]. This substance is involved in a myriad of vital processes, such as secretion, absorption, and transport of macromolecules (e.g. nutrients, hormones, metabolites, antibodies, and neurotransmitters), electrolyte homeostasis, transmission of light and sound, and thermoregulation [2–4]. Therefore, water intake is essential for animal health and survival, particularly in the case of terrestrial vertebrates [3, 5–7].

In all terrestrial mammals, water inputs come from three major sources – water ingested within consumed foods, metabolic water resulting from macronutrient oxidation, and water drunk. Water outputs result from excretion, egestion, or evaporation through the skin or the respiratory tract [4, 5, 8]. When water intake is appropriate, healthy animals maintain a physiological state in which water inputs and outputs are the same throughout the day (i.e. the ‘water balance’), an essential condition for the correct functioning of body cells [2, 5]. Animals reach this water and electrolyte homeostasis by applying a repertoire of behavioral and physiological strategies that depend on the organism’s complexity and the surrounding environment [2]. Whereas drinking increases water input, shade seeking, low metabolic rates, and the excretion of salt by the kidney reduce water loss [1, 2, 4]. Dehydration (i.e. a negative water balance) resulting from long periods of adverse dry conditions when water losses exceed water intake can seriously compromise health, being lethal when losses reach 15 to 25% of body weight (camels are an exception [2, 4]).

Given that plant foods contain more water than animal foods, herbivorous mammals are expected to obtain a larger volume of water from their diets than do omnivores and carnivores [8]. However, plant items can show wide intraspecific and seasonal variations in chemical composition that influence their importance and reliability as water sources, thereby influencing the animals’ need to drink [9]. Herbivorous mammals inhabiting dry environments, such as desert rodents and camelids, can reach water balance by relying on preformed (i.e. water in plant items) and oxidation (i.e. metabolic water resulting from macronutrient oxidation) water during dry periods [2, 10]. In addition to these water sources, animals inhabiting wetter environments also rely on another major source, drinking water [2, 7, 11]. Drinking is rare (e.g. giraffe, *Giraffa camelopardalis*) or presumably nonexistent in mammals that rely on succulent diets [2]. Arboreal folivores once believed to obtain all their water demands from food have been reported to drink either in captivity (sloth, *Choloepus hoffmanii* [12]) or in the wild (koala, *Phascolarctos cinereus* [13, 14]).

While ground-living species drink water from rarely-depletable sources (e.g. rivers, streams, and lagoons), highly arboreal mammals depend on depletable arboreal reservoirs, such as bromeliads and treeholes (primates [15–18]), or on short lasting rain water on tree branches and leaves (koalas [14], sloths [19]). However, the exploitation of terrestrial water reservoirs by these mammals tends to be rare because their vulnerability to predators likely increases when they descend to the forest floor, as has been observed for other tropical primates [15, 20–23].

Among the highly arboreal Neotropical primates, reports of drinking are restricted to a few social groups of the better-studied taxa, including howler monkeys (*Alouatta* spp.[6, 15, 17, 22, 24–26], spider monkeys (*Ateles geoffroyi* [27]), capuchin monkeys (*Cebus capucinus* [28], *Sapajus libidinosus* [29]), and marmosets (*Callithrix flaviceps* [11]). These monkeys meet their water needs primarily via preformed water [15, 30], although they also drink from arboreal reservoirs or, to a lesser extent, terrestrial sources [15–17, 20].

Two main non-exclusive hypotheses have been proposed to explain the drinking behavior of howler monkeys. The thermoregulatory/dehydration-avoidance hypothesis (TDH) relates drinking to a behavioral strategy for maintaining a positive water balance during the hottest and driest periods of the year [6, 17, 26]. The metabolite detoxification hypothesis (MDH) states that the consumption of large amounts of some plant parts (e.g. mature leaves, branches, and seeds) containing digestion inhibitors (fiber and secondary metabolites) ‘forces’ monkeys to drink more to help in their processing [15, 17, 20, 26]. The trend of anti-herbivory metabolites to increase in plants with increasing latitude [31] further supports the potential relevance of the MDH to howler monkeys living in southern latitudes (e.g. *Alouatta guariba clamitans* and *A. caraya).* The bacterial fermentation of the leaf-rich diet of howlers also requires an appropriate water supply [32], as does the excretion of the higher salt content of leaves [8].

Howlers’ low rates of digestion [32] together with the cumulative water loss via urine, lung evaporation, and sweat over the course of an activity period (i.e. daytime), especially during more active and hot times, and under low air humidity [33], can increase plasma osmolarity and cell dehydration to levels that cause thirst and create circadian rhythms of drinking [2, 30]. Similar drinking rhythms associated with feeding are found in squirrel monkeys (*Saimiri* sp. [34]) and owl monkeys (*Aotus* sp. [35]). Finally, forests inhabited by howler monkeys also show seasonal and site-related differences in thermal environment [36], food availability [6, 37], and the presence and reliability of water sources. Therefore, it is important to identify the factors that modulate their drinking behavior to better understand how habitat patch size and spatial restriction resulting from land use changes can affect their health and survival.

Here we investigate the drinking behavior in wild groups of brown howler monkeys (*A. guariba clamitans*) inhabiting Atlantic Forest fragments in southern Brazil as models of folivorous-frugivorous arboreal mammals. Specifically, we assess (i) the arboreal and terrestrial water sources that these monkeys exploit and how they drink, (ii) the daily frequency and seasonal distribution of drinking records, and (iii) the influence of fragment size, season, ambient temperature, rainfall, and the contribution of fruits, leaves, and flowers to the diet on drinking. We predicted that brown howlers would complement the preformed water obtained from their diet with water from arboreal and terrestrial reservoirs, if available, because the availability of fleshy fruits, flowers and young leaves vary seasonally in the study region [37]. We also predicted a within-day increase in drinking in the afternoon in response to an increase in water demands resulting from higher ambient temperatures and the daily water loss via digestion, excretion, breathing, and sweating [1, 8]. Finally, we predicted that diet composition, climatic variables, and fragment size influence the frequency of drinking. While the TDH will receive support if ambient temperature and rainfall are good predictors of the frequency of drinking, a positive influence of leaf ingestion on water consumption will support the MDH.

## Methods

This investigation followed the ethical guidelines of the International Primatological Society and the legal requirements established by the Ethical Committee of the Zoological Society of London for research with nonhuman primates. All studies met all Brazilian animal care policies and were strictly observational. Furthermore, studies conducted from 2011 to 2019 were approved by the Scientific Committee of the Faculty of Biosciences of the Pontifical Catholic University of Rio Grande do Sul (projects SIPESQ #5933 and 7843).

### Study fragments and groups

We collected data on 14 groups of wild brown howlers inhabiting Atlantic Forest fragments ranging from 1 to 977 ha in the municipalities of Porto Alegre, Viamão, and Santa Maria in the state of Rio Grande do Sul, southern Brazil (Table 1, Fig 1). We classified the fragments in three size categories: small (<1 to 10 ha), medium (>10 to 100 ha) and large (>100 to 1,000 ha; *sensu* [38]). Small and medium fragments in Porto Alegre (S1-S3 and M1) and Viamão (S4-S6; Figure 1) were surrounded by anthropogenic matrices comprised of small human settlements, pastures, subsistence orchards, and small parcels of cultivated land (<0.5 to 2 ha). None of them are officially protected. Conversely, the large fragments in Porto Alegre and Viamão (L1-L3) are found in legally protected areas (Fig 1, see [37] for further information on these fragments). The Atlantic Forest study fragments in Santa Maria (S7, M2, L4, and L5; <1 to 977 ha; Figure 1) compose a 5,876-ha mosaic of natural grasslands, extensive pastures devoted to cattle ranching, and other scattered forest fragments. This area, named Campo de Instrução de Santa Maria (CISM), belongs to the Brazilian Army. Therefore, although it is not officially protected by Brazilian laws, CISM is impacted by a lower human pressure than the unprotected study sites.

**Fig 1.**
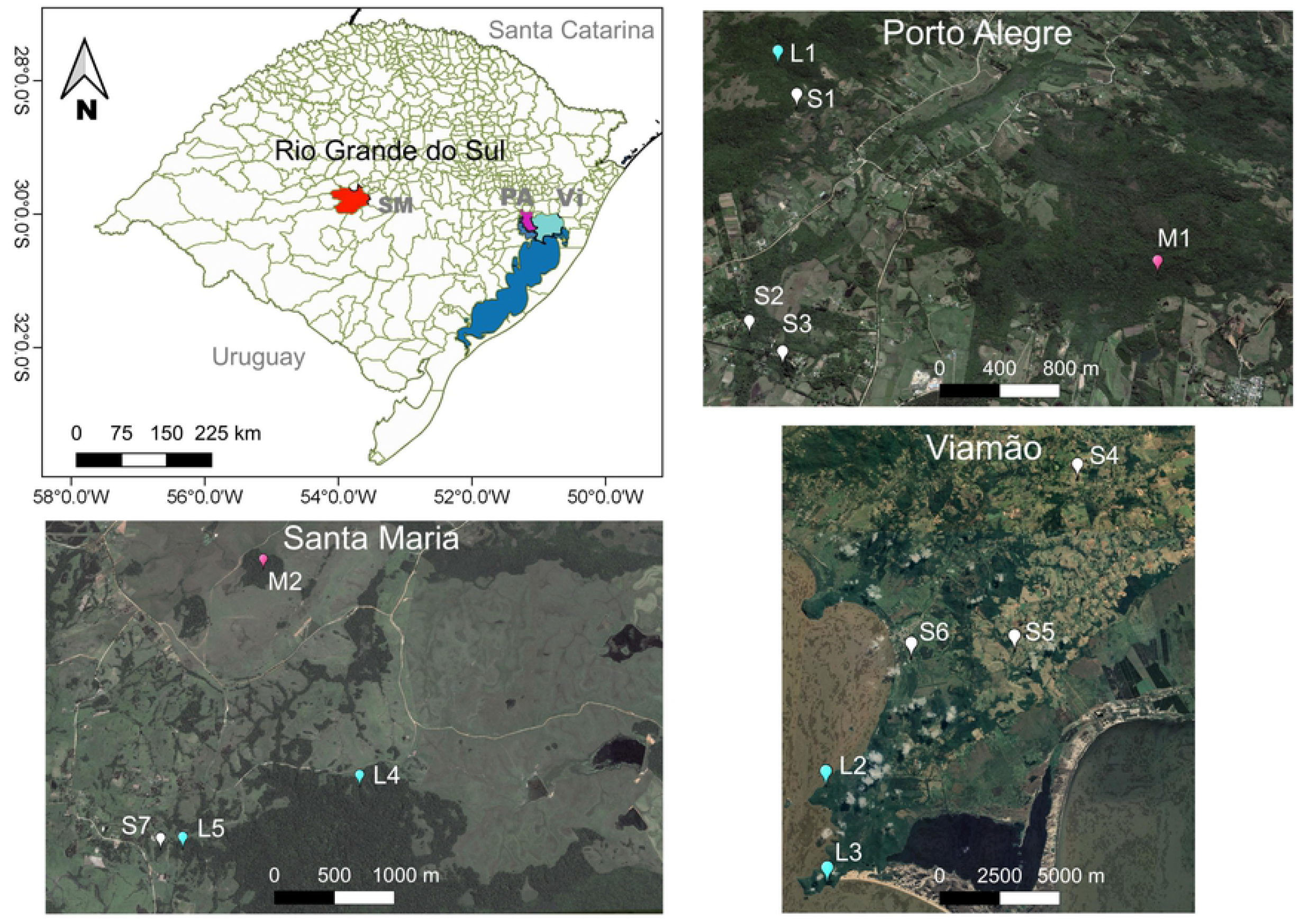
Location of the 14 study sites in the municipalities of Santa Maria (SM, red polygon), Porto Alegre (PA, purple polygon), and Viamão (Vi, cyan polygon), southern Brazil. Color markers indicate the exact location of the small (white), medium (rose), and large (cyan) Atlantic Forest fragments inhabited by the study groups. Lansat7 open-access images (available at http://earthexplorer.usgs.gov/) from 2008 for SM and 2013 for PA and Vi.

**Table 1.** Study fragments, brown howler group size, sampling effort, and number of drinking records.

| Site | Size | Latitude | Longitude | WS<sup>a</sup> | Group size<sup>b</sup> | Sampling effort |  |  | #rec.<sup>d</sup> |
| --- | --- | --- | --- | --- | --- | --- | --- | --- | --- |
|  |  |  |  |  |  | Months | Days<sup>c</sup> | Hours |  |
| S1 | 1.6 | S30°11'00.1" | W51°06'06.6" | P,B | 7 (1,2) | 21 | 67 (4) | 492 | 4 |
| S2 | 9.5 | S30°12'18.4" | W51°06'05.7" | R,B | 11 (1,3) | 19 | 61 (13) | 438 | 47 |
| S3 | 2.3 | S30°12'26.6" | W51°05'54.0" | S,P,B | 10 (1,3) | 20 | 65 (12) | 539 | 28 |
| S4 | 3.6 | S30°12'27.0" | W50°55'39.0" | B,T | 8 (1,2) | 12 | 56 (33) | 681 | 90 |
| S5 | 5.2 | S30°17'27.0" | W50°57'36.0" | B,T | 4 (1,2) | 12 | 55 (14) | 663 | 27 |
| S6 | 7.3 | S30°17'39.0" | W51°00'42.0" | B,T | 8 (1,2) | 12 | 69 (9) | 826 | 13 |
| S7 | 1 | S29°47'05.8" | W53°53'12.0" | S,B | 7 (1,4) | 12 | 59 (44) | 654 | 322 |
| M1 | 27 | S30°12'00.0" | W51°04'00.0" | P,B | 12 (2,2) | 12 | 57 (34) | 518 | 173 |
| M2 | 17 | S29°45'21.3" | W53°52'32.2" | S,B | 6 (1,2) | 12 | 58 (26) | 623 | 99 |
| L1 | 108 | S30°10'39.5" | W51°06'18.2" | R,B,S | 9 (2,3) | 21 | 73 (6) | 536 | 18 |
| L2 | 93 | S30°23'15.6" | W51°02'43.3" | L,P,B | 12 (3,3) | 18 | 81 (7) | 460 | 10 |
| L3 | 106 | S30°20'56.8" | W51°02'58.2" | L,P,B | 10 (2,3) | 17 | 87 (27) | 536 | 102 |
| L4 | 977 | S29°46'46.0" | W53°51'52.0" | S,B | 5 (2,3) | 12 | 54 (45) | 577 | 184 |
| L5 | 977 | S29°47'05.9" | W53°53'03.0" | S,B | 7 (2,3) | 12 | 67 (39) | 836 | 144 |
| **Sum** |  |  |  |  | **116** | **212** | **909 (313)** | **8379** | **1261** |

<sup>a</sup> Water sources detected during the study period: bromeliads (B), treeholes (T), pools (P), rivers (R), streams (S), and lagoon (L).
<sup>b</sup> Group size and number of adult males and females (in parentheses).
<sup>c</sup> Number of study days with drinking events in parentheses.
<sup>d</sup> Total number of drinking records per study group.

The predominant vegetation in all study fragments is subtropical semideciduous forest. Given its latitude (30°-31°S), the region is characterized by marked climatic seasonality: summer (21 December-20 March), fall (20 March-21 June), winter (21 June-22 September), and spring (22 September-21 December). According to meteorological records of Porto Alegre, the average annual ambient temperature during the study period was 21°C [39]. The highest temperatures occurred in the summer (mean=26°C, range=19°-35°C), and the lowest in the winter (mean=15°C, range=3°-26°C; Supplementary Figs S1 and S2). The average annual rainfall during the study years was 1,450 mm. There was no clear rainfall pattern between months or seasons in Porto Alegre,Viamão or Santa Maria (Supplementary Figs S1 and S2).

Despite the variation in fragment size, all fragments contained fleshy fruit tree species (i.e. *Ficus* spp., *Eugenia* spp., and *Syagrus romanzoffiana*) intensively exploited by brown howlers [37, 40, 41]. All study fragments had arboreal and terrestrial water reservoirs, such as bromeliads, streams, and/or rivers (Table 1).

We followed brown howler monkey groups ranging from 4 to 12 individuals in each fragment (*n*=116 individuals, Table 1). All groups in small fragments were well-habituated to humans before study, while we habituated the groups inhabiting medium and large fragments during two to three months prior to their respective monitoring. Whereas most groups inhabited a single forest fragment, S1, S2, and S7 (hereafter named by the acronym of their respective fragments) used more fragments. S1 ranged outside of its most used fragment for about 35% of the study days to feed in a neighboring 10-ha fragment. S2 regularly used three forest fragments distant about 70 to 850 m from each other (the home range of this group included the area of these three fragments). Lastly, S7 also used three forest remnants distant from 30 to 40 m from each other.

### Behavioral data collection

We studied the diet and drinking behavior of the groups during periods ranging from 12 to 21 months (Table 1): (i) January to December 1996 (L5), (ii) June 2002 to August 2003 (M1), (iii) January to December 2005 (S7, M2, and L4), (iv) June 2011 to June 2014 (S1, S2, S3, L1, L2, and L3), and (v) June 2018 to July 2019 (S4, S5, and S6). We collected data for all groups from dawn to dusk using high-resolution 10 x 42 binoculars. We monitored the groups on a monthly basis during three to eight consecutive days in periods (i), (ii), and (iii), during four to five consecutive days on a bimonthly basis in period (iv), and during four to eight consecutive days on a monthly basis in period (v). We recorded the behavior of these groups using the instantaneous scan sampling method in periods (i) to (iv) and the focal-animal method [42] n period (v). However, we recorded all drinking events (i.e. when at least one member of the study group drank) that occurred outside scan or focal sampling units using the ‘all occurrences’ method [42] in all groups. We recorded the behavior of adults, subadults, and conspicuous juvenile individuals, except for S4, S5, and S6, of which we only recorded the behavior of adults.

During feeding bouts we recorded the main plant items eaten (i.e. ripe and unripe fruits, old and young leaves, and flowers) and, whenever possible, the plant species (see [37] for additional details). We used the number of drinking records (i.e. the total number of individual records devoted to drinking per study day) and the number of feeding records devoted to each plant item in the analyses.

### Climatic data

We obtained data on ambient temperature and rainfall for Porto Alegre and Santa Maria from the meteorological database of the Instituto Nacional de Meteorologia do Brasil [39]. We estimated both the mean ambient temperature and the weekly rainfall (i.e. the rainfall accumulated during the previous seven days) for each day with a record of drinking as they represent better the thermal environment and the amount of rainfall water potentially available for brown howlers. Furthermore, we recorded the ambient temperature in the shade at a height of ca. 2 m above the ground after each behavioral sampling unit using a pocket thermo-hygrometer (Yi Chun®, PTH 338) during period (iv) and a portable meteorological station (Nexus, model 351075) distant about 1 km from the study fragments during period (v).

### Statistical analyses

We performed Chi-square tests for proportions to compare the proportions of drinking records per water source and season in each study group using the ‘prop.test’ function of R. We calculated these proportions by dividing the number of records for each water source (or season) by the total number of records for each group during the entire study period. We did not compare fragments or groups because of their sampling effort differences (i.e. the number of sampling months, days, or days per month varied between the five study periods, Table 1). We used the same procedure above to calculate and compare the proportion of drinking records in each hour of the day in those fragments with >10 drinking records. When we found significant differences, we compared the proportion of records in each class using post-hoc proportion contrasts via the R function ‘pairwise.prop.test’ with a Bonferroni correction because of multiple comparisons of the same data sets.

We performed generalized linear mixed-effects models GLMM to assess the influence of the contribution of fruits, leaves, and flowers to the diet, fragment size, ambient temperature, and weekly rainfall on the daily number of drinking records (our response variable) using the function ‘lmer’ of the R package lme4. We set the Poisson error family for the response variable and we specified group ID as a random factor to account for repeated-measures from the same groups. We did not consider interactions between predictor variables to minimize overparameterization and problems of convergence of the global model (i.e. the model containing all fixed and random factors [43]). We standardized variable scales using the ‘stdize’ function of the R package MuMIn [44]. Additionally, we found no multicollinearity problem between variables using the ‘vifstep’ function of R package dplyr [45], as all of them had Variance Inflation Factor (VIF) <3 [46]. Therefore, we included all variables in the global GLMM model.

We used the Akaike’s Information Criterion for small samples (AICc) to select the models that best explain the effects of the predictor variables on drinking behavior. According to this criterion, the model with the strongest empirical support is the one with the smallest difference in AICc [47]. However, given that all models with ΔAICc<2 are considered equally parsimonious, we used the full-model averaging framework to determine which parameters best predict the number of drinking records while accounting for model uncertainty [43]. We used the ‘dredge’ function of the package MuMIn [44] to generate a full submodel set from the global model and the ‘model.avg’ function of the same package to determine the averaged model and the relative importance of each variable or predictor weight (∑*w_i_*). We used a likelihood ratio test over the function ‘anova’ to test the significance of the averaged model compared with the model including only the random factor (i.e. null model). We used the ‘r.squaredGLMM’ function of the package MuMIn to estimate an equivalent of the coefficient of determination or pseudo-*R*^2^ for each competing best GLMM model. All statistical analyses were run in R v.3.6.3 [48] and the statistical significance threshold was set at *P*≤0.05.

## Results

### Water sources

We obtained a total of 1,258 individual drinking records (range=4-322 records/group, Table 1) distributed in 917 events of group drinking and 313 observation days (range=0-16 records/day, Table 1). We did not record drinking in 66% of the study days (i.e. 596 out of 909 days).

The water sources were streams (44% of 1,258 records), followed by treeholes (26%), *Vriesea*, *Aechmea*, and *Tillandsia* bromeliads (16%), pools (11%), and rivers (3%) (Fig 2a). The proportion of drinking records per water source type differed in nine of the fourteen groups (*X*^2^ tests, *P*<0.05 in all significant cases, Fig 2a). Arboreal sources were exploited by most groups (treeholes=12, bromeliads=11), whereas terrestrial ones were less common (streams=6, pools=4, rivers=2; Fig 2a).

**Fig 2.**
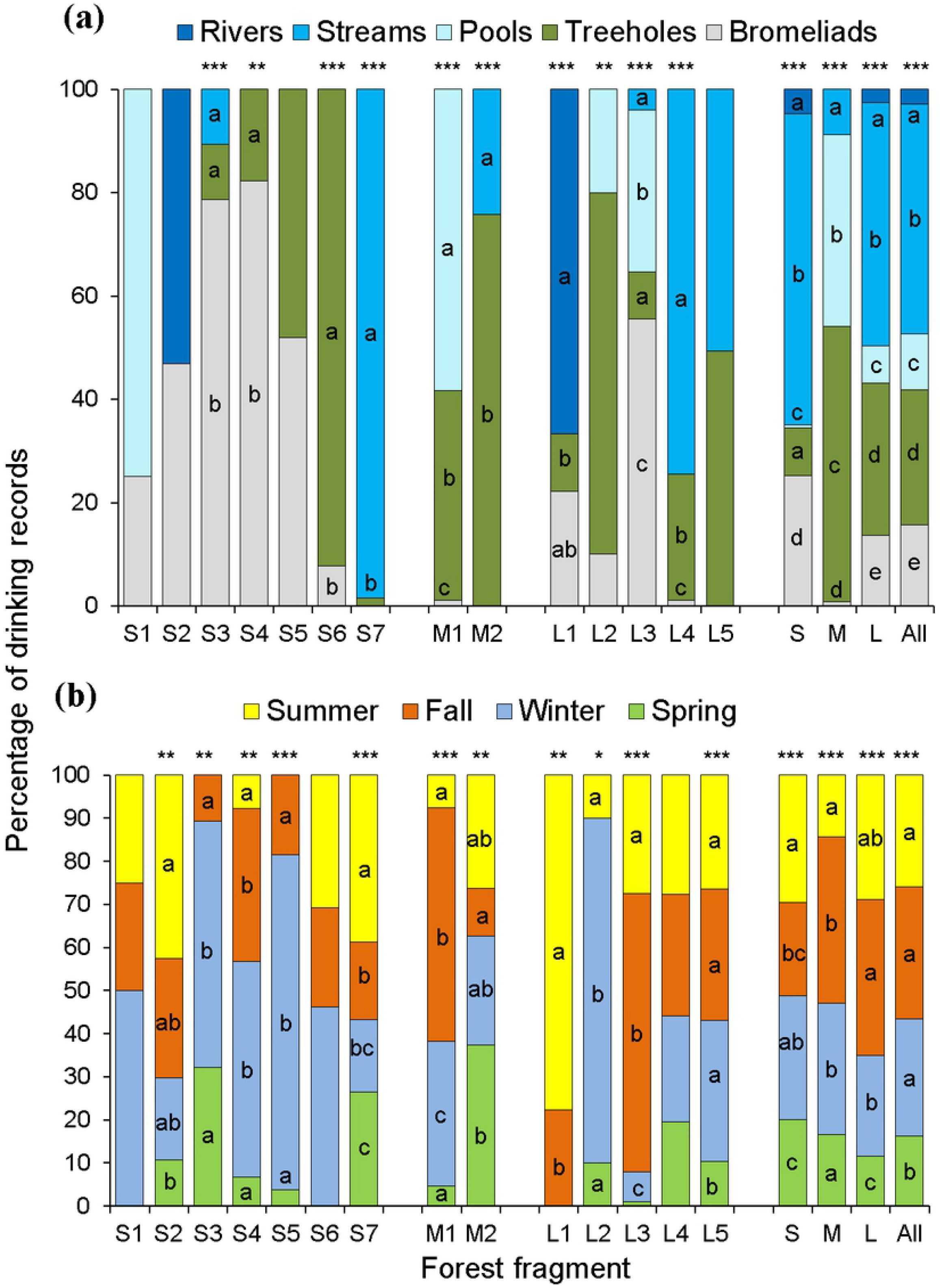
Percentage of drinking records in 14 brown howler groups per water source (a) and season (b). The number of records is indicated in the middle of each bar. Asterisks on the bars indicate the significance level according to Chi-square tests for proportions: *\*P≤0.05,* \*\**P*<0.01, \*\*\**P*<0.001. The proportion of records at the forest size level (small=S, medium=M, and large=L) and in the pooled dataset (All) is indicated in the four bars to the right. Water sources – rivers: permanent water currents >4-m in width and >1-m in depth; streams: seasonal water currents <2-m in width and <1-m in depth; treeholes: 10-40-cm diameter holes in trunks or large branches; bromeliads: water stored in the rosette of epiphytic bromeliads. Significant differences in the proportion of records between water sources or seasons within each fragment are indicated with different lower-case letters in the bars. When proportion contrasts tests did not detect differences, no letter is shown. *N*=1,128 records in (a) and 1,131 records in (b).

The most common drinking behavior consisted of inserting their head and sipping water directly from bromeliads and treeholes. When the treehole had a small diameter, monkeys immersed a cupped hand into the hole, pulled it out, and placed the mouth under the fingers to lick the dripping water. Vigilance was negligible during these arboreal drinking events.

In contrast, when drinking from terrestrial sources (rivers, streams and pools) howlers scanned the surroundings very carefully and were highly vigilant when drinking. Terrestrial drinking events began with some group members moving slowly to the understory, where they remained vigilant for ca. 30 s to 5 min before one or two of them descended to the ground to drink directly from the terrestrial water source for 102 ± 66 s (mean ± S.D., *n*=463) while the other individuals waited in vigilance in the understory. When the first individuals climbed back to the understory, the others descended slowly to the ground to drink, and the first remained in vigilance. A single drinking event involved between 1/5 and 4/5 of the group members.

### Seasonal and daily patterns in drinking behavior

We found no clear pattern in the proportion of drinking records between or within seasons (Fig 2b, Supplementary Fig S3). We observed drinking in all seasons in seven fragments, in three seasons in six fragments, and in two seasons in the remaining fragment (Fig 2b). For those fragments where we found seasonal differences in the proportion of drinking records (*n*=11, proportion contrasts, *P*<0.05 in all significant cases, Fig 2b), a greater proportion of records occurred in a single season (winter - *n*=3 fragments: S3, S5, and L2; summer - *n*=2 fragments: S7 and L1; fall - *n*=2 fragments: M1, L3), in two seasons (*n*=1 fragment: S4) or in three (*n*=3 fragments: S2, M2, and L5; Fig 2b). We found a lower percentage of drinking records in the spring than in the other seasons in the pooled dataset (*X*^2^ =77, df=3, *P*<0.0001; Fig 2b).

Finally, we found that the distribution of drinking during the day showed a unimodal pattern in most fragments. The higher percentages of records occurred in the afternoon, particularly from 15:00 to 17:00 (8 out of 12 analyzed fragments; proportion contrasts, *P*<0.05 in all significant cases, Fig 3). This peak of drinking occurred near times with higher ambient temperatures in the fragments for which we have in-site temperature data (*n*=7; Supplementary Fig S4).

**Fig 3.**
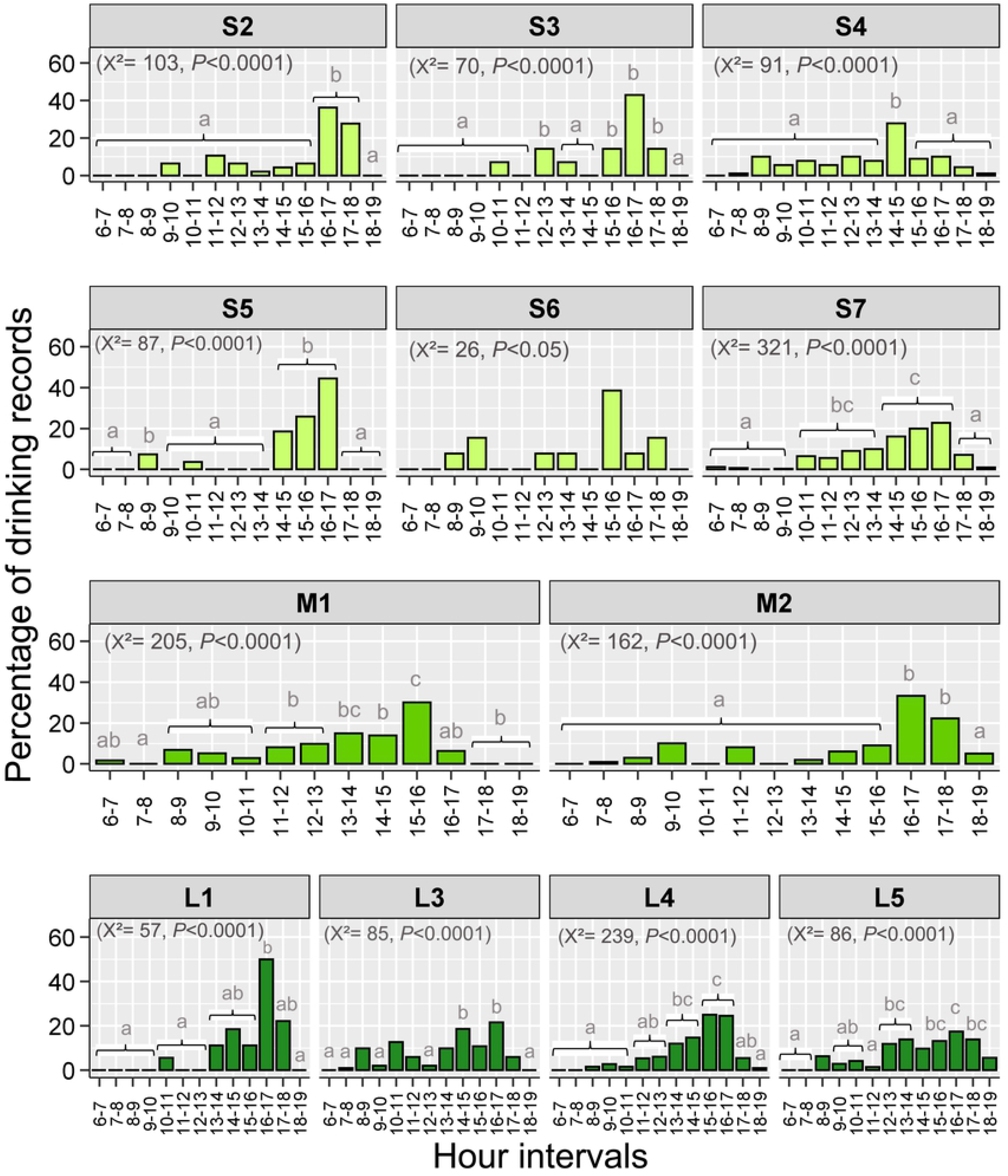
Variation in percentage of drinking records by brown howler monkeys during the day in small, medium, and large Atlantic Forest fragments. Different lower-case letters on the bars indicate significant differences after Bonferroni adjustment in *P* values. The absence of letters indicates that these hour intervals did not differ from the others. The number of degrees of freedom was 12 in all cases. Only fragments with >10 drinking records were considered in this analysis.

### Factors driving the drinking behavior of brown howlers

We found six models that included all predictor variables, except weekly rainfall, with substantial empirical support (i.e. ΔAICc<2; Table 2). Flower and leaf consumption were the only predictors present in all models. The two best models for explaining the frequency of drinking contained only these two variables (first), plus ambient temperature (second; Table 2). The averaged model differed from the null model (likelihood ratio test: *X*^2^=22, df=5, *P*<0.001). Flower consumption had an inverse relationship with drinking (β=-0.14, *z-*value=3.08, *P*<0.01), whereas leaf consumption had a direct one (β= 0.14, *z-*value=2.35, *P*<0.05, Table 2). Fragment size, fruit consumption, and ambient temperature had insignificant relationships with howler monkey drinking (Table 2).

**Table 2.** Best supported GLMM models (ΔAICc<2) and model-averaged that predict the variation in the number of drinking records in 14 brown howler groups in southern Brazil.

| Predictor variables<sup>a</sup> | Parameters<sup>b</sup> |  |  |  |  |
| --- | --- | --- | --- | --- | --- |
| *Best supported models* |  |  |  |  |  |
|  | AIC<sub>c</sub> | ΔAIC<sub>c</sub> | w<sub>i</sub> | R<sup>2</sup><sub>c</sub> |  |
| 1) Flower+Leaf | 2049.6 | 0.00 | 0.23 | 0.53 |  |
| 2) Flower+Leaf+Temp | 2049.7 | 0.04 | 0.23 | 0.53 |  |
| 3) Flower+Fsize+Leaf | 2050.2 | 0.54 | 0.18 | 0.54 |  |
| 4) Flower+Fsize+Leaf+Temp | 2050.2 | 0.59 | 0.17 | 0.54 |  |
| 5) Flower+Fruit+Leaf | 2051.3 | 1.63 | 0.10 | 0.54 |  |
| 6) FLower+Fsize+Leaf+Temp | 2051.5 | 1.86 | 0.09 | 0.54 |  |
| *Averaged model (R<sup>2</sup><sub>c</sub>=0.55)* |  |  |  |  |  |
|  | β<sub>i</sub> | SE | 95% CI | z-value | Σw<sub>i</sub> |
| Intercept | 2.03 | 0.1 | (1.8, 2.2) | 19.4*** | — |
| Flower consumption (Flower) | -0.14 | 0.05 | (-0.23, -0.05) | 3.08** | 1 |
| Leaf consumption (Leaf) | 0.14 | 0.06 | (0.02, 0.25) | 2.35* | 0.95 |
| Ambient temperature (Temp) | 0.03 | 0.04 | (-0.02, 0.15) | 0.69 | 0.47 |
| Fragment size (Fsize) | 0.1 | 0.2 | (-0.17, 0.77) | 0.52 | 0.42 |
| Fruit consumption (Fruit) | 0.01 | 0.03 | (-0.08, 0.15) | 0.23 | 0.26 |

<sup>a</sup>Number of parameters in each model ( $k$ ), Akaike's Information Criterion for small samples ( $AIC_c$ ), difference in $AIC_c$ ( $\Delta AIC_c$ ), model probability Akaike weights ( $w_i$ ), Pseudo- $R^2$ ( $R^2_c$ ) indicating the percent of variance explained by the fixed and random factors, partial regression coefficients of the model-averaged ( $\beta$ ), standard errors which incorporate model uncertainty (SE), 95% confidence intervals for the parameter estimates (95% CI), and relative importance of each predictor variable ( $\sum w_i$ ). Significance level: \* $P < 0.05$ , \*\* $P < 0.01$ , \*\*\* $P < 0.001$ .

## Discussion

We found that brown howlers drank water accumulated in bromeliads and treeholes in the canopy, and that they also descended to the ground to drink from streams, rivers, and pools. Drinking increased in the afternoon and was less frequent in the spring. Also, while howlers drank more when their diet included more leaves and drank less when they ate more flowers, the contribution of fruits to the diet, habitat size, mean ambient temperature, and rainfall did not predict the frequency of drinking.

The exploitation of non-food arboreal and terrestrial water reservoirs supports our expectation that oxidation and preformed water are insufficient for permanently satisfying howlers’ water needs, as reported for many terrestrial mammals [1–3]. The finding that streams were the most used water sources by brown howlers differs from the greater importance of arboreal water reservoirs for other howler monkeys inhabiting both large (e.g. 1,564 ha in Barro Colorado Island, Panama [15, 30]) and small forest remnants (e.g. ≤10 ha [26, 49]).

This use of terrestrial water sources occurred despite the high risk of predation by domestic/stray dogs and small wild felids in the study region (e.g. *Leopardus wiedii* [50]; also OM Chaves, personal communication). The fact that dog attacks represent a major cause of brown howler death in urban and suburban populations in southern Brazil [23, 51] explains the highly cautious behavior and vigilance displayed by brown howlers when descending to the ground to drink, a behavior also observed in other primates (e.g. *Callithrix flaviceps* [11]). This threat is believed to reduce (or even eliminate) howlers’ use of terrestrial water reservoirs in better-conserved large forests inhabited by wild carnivore populations in Central America (e.g. *A. palliata* [30, 52]). The frequency of brown howler remains in ocelot (*Leopardus pardalis*) scats in a ca. 950-ha Atlantic Forest reserve in southeastern Brazil highlights their vulnerability to wild felids [21].

The general lower drinking in the spring may be explained, at least partially, by three main reasons. First, unlike at lower tropical latitudes where the hottest and driest times often coincide (i.e. dry season [53]), summer and spring are the hottest, but not the driest seasons in the subtropical study region (ca. 31°S) (Supplementary Fig S1). In fact, rainfall is relatively well distributed throughout the year in Rio Grande do Sul state ([39], see also Supplementary Figs S1 and S2), where ‘rainy quarters’ occur at any time [54].

Second, the higher availability and consumption of flowers and fleshy ripe fruits, and the consequent lower consumption of leaves, by the study groups also occurred in the hottest seasons [37]. This diet composition likely reduces the need for water to detoxify secondary metabolites while supplying water to counterbalance the losses of thermoregulation and other physiological processes.

Third, brown howlers may lower the thermoregulatory demands for water by preventing body over-heating and dehydration via riparian microhabitat selection (OM Chaves and VB Fortes, personal observation), positional adjustments and shade-seeking [55] during the hottest times of the day (strategies also reported in other Neotropical primates: *A. palliata* [56], *A. caraya* [36], *Cebus capucinus* [57], *Callicebus bernhardi* [58]). Despite these strategies, the peaks of drinking in the afternoon tended to occur around the warmer times of the day (Supplementary Fig S4), which are likely triggered by the need of water in this period of intensified physiological thermoregulation together with the recovery of the water spent earlier in the day that is required to reach the homeostasis of blood osmolarity [34, 35]. Testing this hypothesis requires data on body temperature and water balance.

The within-day relationship between drinking and higher ambient temperatures together with the lack of a significant relationship between mean ambient temperature and drinking at a broader temporal scale show that the circadian rhythm of drinking supports the thermoregulatory/dehydration-avoidance hypothesis (TDH), whereas the seasonal pattern of drinking does not. This finding is not surprising given the everlasting essential role that water plays in the functioning of living organisms and the absence of a dry season in the study region. In seasonal environments where water availability decreases significantly during the dry season, howler (*Alouatta palliata*), spider (*Ateles geoffroyi*), and capuchin (*Cebus capucinus*) monkeys may camp around the remaining arboreal and ground water reservoirs [52]. Yet the opposing influences of the consumption of leaves and flowers on the frequency of drinking support the metabolite detoxification hypothesis (MDH). While the ingestion of leafy material can also demand water for the process of bacterial fermentation [30, 32], flowers can have high water contents [59] that contribute to satisfy the monkeys’ daily requirements [60].

In sum, we have found that both the TDH and the MDH can explain the drinking behavior of brown howlers in response to short-term thermal environment and diet composition. Extrapolating from brown howlers to arboreal folivorous-frugivorous mammals in general that also lack adaptations to tolerate high levels of dehydration, we suggest that the higher the proportion of leaves in their diet, the greater might be the challenges in fulfilling their water requirements, particularly in habitats where terrestrial water reservoirs are scarce or absent, such as some forest fragments. Therefore, highly folivorous species may be more sensitive to droughts than more frugivorous ones. Despite the higher availability of leaves than flowers and fruits in forests, highly folivorous mammals may also be more vulnerable to predators if they are forced to descend to the ground to drink from terrestrial reservoirs, particularly in forest fragments immersed in anthropogenic landscapes where dogs roam freely. In this respect, studies assessing how differences in land-use and human disturbance influence the abundance and distribution of arboreal and terrestrial water reservoirs and how they impact the drinking behavior, water balance, and health of arboreal folivorous-frugivorous mammals are critical for enabling us to design and implement appropriate management strategies for promoting their conservation in anthropogenic fragmented landscapes.

## Supporting information

**Fig. S1.** Rainfall (blue line) and mean air temperature (yellow line) in Porto Alegre and Viamão municipallities during the study.

**Fig. S2.** Rainfall (blue) and mean air temperature (yellow) in Santa Maria municipality during the study years of 1996 and 2005.

**Fig. S3.** Seasonal distribution of drinking events by 14 brown howler monkey groups inhabiting small, medium, and large Atlantic forest fragments in southern Brazil.

**Fig. S4.** Hourly variation in average ambient temperature of seven Atlantic Forest fragments in Porto Alegre and Viamäo municipalities.

## Acknowledgements

We thank Danielle Camaratta and João Claudio Godoy for logistical support and field assistance. We thank the landowners of the study fragments in Porto Alegre and Viamão for giving us permission to conduct this research on their properties. We thank Commandant Aluísio S.R. Filho for giving us permission to work in the Campo de Instrução de Santa Maria (CISM).

## Author Contributions

**Conceptualization:** Óscar M. Chaves, Júlio César Bicca-Marques.

**Data curation:** Óscar M. Chaves, Vanessa B. Fortes, Gabriela P. Hass.

**Formal analysis:** Óscar M. Chaves, Júlio César Bicca-Marques.

**Visualization:** Óscar M. Chaves, Júlio César Bicca-Marques.

**Investigation:** Óscar M. Chaves, Júlio César Bicca-Marques, Vanessa B. Fortes, Gabriela P. Hass, Renata B. Azevedo.

**Methodology:** Óscar M. Chaves, Júlio César Bicca-Marques, Vanessa B. Fortes, Gabriela P. Hass, Kathryn E. Stoner.

**Project administration:** Óscar M. Chaves

**Writing-original draft:** Óscar M. Chaves

**Writing-review & editing:** Óscar M. Chaves, Júlio César Bicca-Marques, Vanessa B. Fortes, Gabriela P. Hass, Kathryn E. Stoner.

**Funding acquisition:** Júlio César Bicca-Marques, Kathryn E. Stoner.

## References

1. Nagy KA, Peterson CC. Scaling of water flux rate in animals. London: University of California Press, 1988.

2. Fitzsimons J. Evolution of physiological and behavioural mechanisms in vertebrate body fluid homeostasis. In: Ramsay DJ, Booth D, editors. Thirst. New York: Springer, 1991. pp. 3–22.

3. NRC. Nutrient requirements of nonhuman primates. Washington DC: National Resource Council, National Academies Press, 2003.

4. Jéquier E, Constant F. Water as an essential nutrient: the physiological basis of hydration. Eur J Clin Nutr. 2010; 64:115–123.

5. Grandjean AC, Campbell SM. Hydration: fluids for fife. A monograph by the North American Branch of the International Life Science Institute. Washington DC: ILSI North America, 2004.

6. Dias PA, Rangel-Negrin A, Coyohua-Fuentes A, Canales-Espinosa D. Factors affecting the drinking behavior of black howler monkeys (*Alouatta pigra*). Primates. 2014; 55:1–5.

7. Sharma N, Huffman MA, Gupta S, Nautiyal H, Mendonca R, Morino L, Sinha A. Watering holes: the use of arboreal sources of drinking water by Old World monkeys and apes. Behav Proc. 2016; 129:18–26.

8. McNab BK. The physiological ecology of vertebrates: a view from energetics. London: Cornell University Press, 2002.

9. Green B. Field energetics and water fluxes in marsupials. In: Saunders NR, Hinds LA, editors. Marsupial Biology: recent research, new perspectives. Sydney: UNSW Press, 1997. pp. 143–159.

10. Xu, M.M., Wang, D.H. Water deprivation up-regulates urine osmolality and renal aquaporin 2 in Mongolian gerbils (*Meriones unguiculatus*). Comp Biochem and Phys A. 2016; 194:37–44.

11. Ferrari SF, Hilário RR. Use of water sources by buffy-headed marmosets (*Callithrix flaviceps*) at two sites in the Brazilian Atlantic forest. Primates. 2012; 53:65–70.

12. Gilmore DP, Da-Costa CP, Duarte DPF. An update on the physiology of two- and three-toed sloths. Braz J Med Biol Res. 2000; 33:129–146.

13. Ellis W, Melzer A, Clifton ID, Carrick F. Climate change and the koala *Phascolarctos cinereus:* water and energy. Aust Zool. 2010; 35:369–377.

14. Mella VSA, Orr C, Hall L, Velasco S, Madani G. An insight into natural koala drinking behavior. Ethology. 2020. https://onlinelibrary.wiley.com/doi/abs/10.1111/eth.13032.

15. Glander KE. Drinking from arboreal water sources by mantled howling monkeys (*Alouatta palliata-Gray*). Folia Primatol. 1978a; 29:206–217.

16. Oates JF. Water-plant and soil consumption by guereza monkeys (*Colobus guereza):* a relationship with minerals and toxins in the diet? Biotropica. 1978; 10:241–253.

17. Bicca-Marques JC. Drinking behavior in the black howler monkey (*Alouatta caraya*). Folia Primatol. 1992; 58:107–111.

18. Hillyer AP, Armstrong R, Korstjens AH. Dry season drinking from terrestrial man-made watering holes in arboreal wild Temminck’s red colobus, the Gambia. Primate Biol. 2015; 2:21–24.

19. Martínez N, Antelo C, Rumiz DE. Rehabilitación de perezosos (*Bradypus variegatus*) urbanos en reservas privadas aledañas a Santa Cruz de La Sierra: una iniciativa multipropósito de investigación, manejo y educación. Rev Bol Ecol. 2004; 16:1–10.

20. Glander KE. Howling monkey feeding behavior and plant secondary compounds: a study of strategies. In: Montgomery GG, editor. The ecology of arboreal folivores, Washington DC: Smithsonian Institution Press, 1978b. pp. 561–574.

21. Bianchi RC, Mendes SL. Ocelot (*Leopardus pardalis*) predation on primates in Caratinga Biological Station, southeast Brazil. Am J Primatol. 2007; 69:1173–1178.

22. Pozo-Montuy G, Serio-Silva JC. Movement and resource use by a group of *Alouatta pigra* in a forest fragment in Balancán, México. Primates. 2007; 48:102–107.

23. Bicca-Marques JC, Chaves ÓM, Hass GP. Howler monkey tolerance to habitat shrinking: lifetime warranty or death sentence? Am J Primatol. 2020; 82:e23089.

24. Bonvicino CR. Observações sobre a ecologia e o comportamento de *Alouatta belzebul* (Primates: Cebidae) na Mata Atlântica. M.Sc.Thesis, Universidade Federal da Paraíba. 1988.

25. Steinmetz S. Drinking by howler monkeys (*Alouatta fusca*) and its seasonality at the Intervales State Park, São Paulo, Brazil. Neotrop Primates. 2001; 9:111–112.

26. Moro-Rios RF, Serur-Santos CS, Miranda JMD, Passos FC. Water intake by a group of *Alouatta clamitans* (Primates: Atelidae), on an Araucaria Pine Forest: seasonal, sex-age and circadian variations. Rev Bras Zool. 2008; 25:558–562.

27. Campbell CJ, Aureli F, Chapman CA, Ramos-Fernandez G, Matthews K, Russo SE, Suarez S, Vick L. Terrestrial behavior of *Ateles* spp. Int J Primatol. 2005; 26:1039–1051.

28. Freese CH. The behavior of white-faced capucins (*Cebus capucinus*) at a dry-season waterhole. Primates. 1978; 19:275–286.

29. Castro SCN, Souto AS, Schiel N, Biondi LM, Caselli CB. Techniques used by bearded capuchin monkeys (*Sapajus libidinosus*) to access water in a semi-arid environment of North-Eastern Brazil. Folia Primatol. 2017; 88:267–273.

30. Nagy KA, Milton K. Aspects of dietary quality, nutrient assimilation and water-balance in wild howler monkeys (*Alouatta palliata*). Oecologia. 1979; 39:249–258.

31. Moles AT, Bonser SP, Poore AG, Wallis IR, Foley WJ. Assessing the evidence for latitudinal gradients in plant defense and herbivory. Func Ecol. 2011; 25:380–388.

32. Milton K. Food choice and digestive strategies of two sympatric primate species. Am Nat. 1981; 117:476–495.

33. Thompson CL, Williams SH, Glander KE, Teaford MF, Vinyard CJ. Body temperature and thermal environment in a generalized arboreal anthropoid, wild mantled howling monkeys (*Alouatta palliata*). Am J Phys Anthropol. 2014; 154:1–10.

34. Sulzman FM, Fuller CA, Moore-Ede M. Feeding time synchronizes primate circadian rhythms. Physiol Behav. 1977; 18:775–779.

35. Hoban TM, Levine AH, Shane RB, Sulzman FM. Circadian rhythms of drinking and body temperature of the owl monkey (*Aotus trivirgatus*). Physiol Behav. 1985; 34:513–518.

36. Bicca-Marques JC, Calegaro-Marques C. Behavioral thermoregulation in a sexually and developmentally dichromatic neotropical primate, the black-and-gold howling monkey (*Alouatta caraya*). Am J Phys Anthropol. 1998; 106:533–546.

37. Chaves ÓM, Bicca-Marques JC. Feeding strategies of brown howler monkeys in response to variations in food availability. Plos One. 2016; 11:e0145819.

38. Marsh LK, Chapman CA, Norconk MA, Ferrari SF, Gilbert KA, Bicca-Marques JC, Wallis J. Fragmentation: specter of the future or the spirit of conservation? In: Marsh LK, editor. Primates in fragments: ecology and conservation. New York: Kluwer Academic/Plenum Publishers, 2003. pp. 381–398.

39. INMET. Banco de Dados meteorológicos para ensino e pesquisa: registros metereologicos para Rio Grande do Sul. Instituto Nacional de Metereologia do Brasil, Porto Alegre, 2019. https://www.inmet.gov.br/portal/index.php?r=bdmep/bdmep.

40. Chaves ÓM, Bicca-Marques JC. Dietary flexibility of the brown howler monkey throughout its geographic distribution. Am J of Primatol. 2013; 75:16–29.

41. Fortes VB. Ecologia e comportamento do bugio-ruivo (*Alouatta guariba clamitans* Cabrera, 1940) em fragmentos florestais na depressão central do Rio Grande do Sul, Brasil. Ph.D. Thesis, Pontifícia Universidade Católica do Rio Grando do Sul. 2008. http://repositorio.pucrs.br/dspace/bitstream/10923/5421/1/000400984-Texto%2BCompleto-0.pdf.

42. Altmann J. Observational study of behavior: sampling methods. Behavior. 1974; 49:227–267.

43. Grueber CE, Nakagawa S, Laws RJ, Jamieson IG. Multimodel inference in ecology and evolution: challenges and solutions. J Evol Biol. 2011; 24:699–711.

44. Barton K. MuMIn: multi-model Inference. R package version 1.15.6. http://cran.r-project.org/web/packages/MuMIn/index.html. 2016.

45. Wickham H, François R, Henry L, Müller K. dplyr: a grammar of data manipulation. R package version 0.7.8. https://CRAN.R-project.org/package=dplyr. 2018.

46. Zuur AF, Ieno EN, Elphick CS. A protocol for data exploration to avoid common statistical problems. Methods Ecol Evol. 2010; 1:3–14.

47. Burnham KP, Anderson D. Model selection and multi-model inference: a practical information-theoretic approach. New York: Springer Press; 2003.

48. R Core Team. R: a language and environment for statistical computing. Version 3.6.3. https://www.R-project.org. 2020.

49. Silver SC, Ostro LET, Yeager CP, Horwich R. Feeding ecology of the black howler monkey (*Alouatta pigra*) in northern Belize. Am J Primatol. 1998; 45:263–279.

50. Gonçalves AS. Uso de hábitat de mamíferos terrestres em fragmentos de floresta estacional decidual. M.Sc. Thesis, Universidade do Vale do Rio dos Sinos. 2006. http://www.repositorio.jesuita.org.br/bitstream/handle/UNISINOS/2300/uso%20de%20habitat.pdf?sequence=1.

51. Buss G. Conservação do bugio-ruivo (*Alouatta guariba clamitans*) (PRIMATES, ATELIDAE) no entorno do Parque Estadual de Itapuã, Viamão, RS. Ph.D. Thesis, Universidade Federal do Rio Grande do Sul. 2012. https://www.lume.ufrgs.br/bitstream/handle/10183/69918/000867969.pdf?sequence=1.

52. Chapman CA. Patterns of foraging and range use by three species of Neotropical primates. Primates. 1988; 29:177–194.

53. Allaby M. Tropical forests. New York: Chelsea House Publishers, 2006.

54. Nimer E. Climatologia do Brasil. 2.ed. Rio de Janeiro, Brazil: IBGE/Departamento de Recursos Naturais e Estudos Ambientais, 1989.

55. Bicca-Marques JC, Azevedo RB. The ‘thermoregulation hypothesis’ does not explain the evolution of sexual dichromatism in the brown howler monkey (*Alouatta guariba clamitans*). Folia Primatol. 2004; 75:236.

56. Paterson JD. Postural-positional thermoregulatory behaviour and ecological factors in primates. Can Rev Phys Anthropol. 1981; 3:3–11.

57. Campos FA, Fedigan LM. Behavioral adaptations to heat stress and water scarcity in white-faced capuchins (*Cebus capucinus*) in Santa Rosa National Park, Costa Rica. Am J Phys Anthropol. 2009; 138:101–111.

58. Lopes KG, Bicca-Marques JC. Ambient temperature and humidity modulate the behavioural thermoregulation of a small arboreal mammal (*Callicebus bernhardi*). J Thermal Biol. 2017; 69:104–109.

59. Richard A. Primates in nature. New York: WH Freeman, 1985.

60. Hogan JD, Melin AD, Mosdossy KN, Fedigan LM. Seasonal importance of flowers to Costa Rican capuchins (*Cebus capucinus imitator*): implications for plant and primate. Am J Phys Anthropol. 2016; 161:591–602.

